# Influenza A virus infection alters lipid packing and surface electrostatic potential of the host plasma membrane

**DOI:** 10.1101/2023.07.25.550511

**Authors:** Annett Petrich, Salvatore Chiantia

## Abstract

The pathogenesis of influenza A viruses (IAVs) is influenced by several factors, including IAV strain origin and reassortment, tissue tropism and host type. While such factors were mostly investigated in the context of virus entry, fusion and replication, little is known about the viral-induced changes to the host lipid membranes which might be relevant in the context of virion assembly. In this work, we applied several biophysical fluorescence microscope techniques (i.e., Förster energy resonance transfer, generalized polarization imaging and scanning fluorescence correlation spectroscopy) to quantify the effect of infection by two IAV strains of different origin on the plasma membrane (PM) of avian and human cell lines. We found that IAV infection affects the membrane charge of the inner leaflet of the PM. Moreover, we showed that IAV infection impacts lipid-lipid interactions by decreasing membrane fluidity and increasing lipid packing. Because of such alterations, diffusive dynamics of membrane-associated proteins are hindered. Taken together, our results indicate that the infection of avian and human cell lines with IAV strains of different origins had similar effects on the biophysical properties of the PM.

## 1. Introduction

Influenza A virus (IAV) is an enveloped, negative-sense RNA virus that belongs to the Orthomyxoviridae family [1, 2]. This pathogen poses a significant threat to both humans and animals and can cause widespread infections resulting in significant morbidity and mortality [2, 3]. Apart from the commonly occurring seasonal IAV subtypes H1N1 and H3N2, there has been a recent rise in human infections caused by different avian influenza viruses such as H5Nx, H7N9, and H9N2, as well as swine influenza viruses [2, 3]. Several studies have revealed that IAV infection can alter the lipid metabolism in various hosts, impacting IAV replication, viral envelope lipid composition and potentially contributing to pathogenicity [1, 4-9]. These alterations have been also linked to inflammatory responses in host organisms [1, 8]. Additionally, some avian IAV strains have been found to trigger a more intense inflammatory response in humans compared to human(-adapted) strains [10-12]. Therefore, investigating the connection between lipid metabolism, physical properties of cellular membranes and IAV infection is crucial for understanding the pathogenic mechanisms of IAV and developing targeted antiviral treatments.

In general, lipids and cellular membranes play a crucial role in various stages of the IAV life cycle, such as virus-host receptor interaction, membrane fusion, nuclear transport, virion assembly and budding [11, 13-17]. Several studies have proposed that IAV assembles and buds from specific lipid domains within the apical plasma membrane (PM), which are enriched in cholesterol and sphingolipids [15, 16, 18-21]. This hypothesis is supported by lipidome analyses conducted on purified influenza viral envelopes, which were shown to contain higher levels of certain sphingolipid species and cholesterol, thus potentially increasing the bilayer structural order in comparison to the host-cell membrane [5, 19, 21, 22]. Furthermore, removing cholesterol from the viral envelope destabilized the viral membrane and morphology, leading to a decrease in virus infectivity [23, 24]. Of note, a previous study found that three egg-grown IAV strains with varying pathogenicity exhibited a modified glycerophospholipid (GPL) composition compared to non-infected allantoic fluid (NAF) or mammalian cells [5]. Specifically, highly pathogenic IAV strains were found to contain higher fractions of sphingomyelin (SM) and saturated fatty acids compared to the other IAV strains and NAF [5].

On the other hand, it is also well established that IAV structural proteins participate in specific lipid-protein interactions in infected cells. For example, the spike proteins hemagglutinin (HA) and neuraminidase (NA) are transported to the PM via cholesterol-/sphingomyelin-rich vesicles [13, 14, 17, 18, 20, 25, 26] and their lateral organization might be indirectly influenced by lipids [27, 28] or specifically depend on ordered lipid-protein domains [20, 29, 30]. The cytoplasmic tail of HA possesses multiple basic residues which interact with phosphoinositides (e.g., PIP2), modulating protein clustering and membrane association [31, 32]. Additionally, several studies have investigated the interplay between lipids and the IAV matrix protein 2 (M2) [30, 33-40]. It has been proposed that M2 is transported to the apical PM independently from HA and NA through its interaction with the phosphatidylserine (PS)-conjugated microtu-bule-associated protein 1 light chain 3 protein (LC3-II) [37, 41-43]. Thus, the interaction between M2 and the IAV matrix protein 1 (M1) might be supported by a local enrichment of PS at the virus assembly site [44-46]. Moreover, previous studies have indicated that M2-M2 interactions are enhanced in cholesterol-enriched membranes [35, 39, 40].

These findings emphasize the crucial role of a controlled lipid metabolism in multiple stages of IAV infection, which appears to be influenced by both the origin of the IAV strain and the host type [1, 4-11, 14, 17, 47]. While previous studies have primarily investigated virus entry, fusion, and replication, a comprehensive analysis of how the host environment and IAV strain origin impact virus assembly and the PM environment in general is still lacking. Previous lipidomic studies focused mainly on the analysis of purified influenza virions [5, 19, 21, 22] and only few analyzed the lipidome of whole (non-)infected mammalian cells [6, 7, 9, 22]. A small number of studies observed no significant difference in the lipidome of whole cells after infection compared to non-infected cells whereas others reported IAV-mediated induction of sphingomyelin, cholesterol and fatty acid biosynthesis [6, 7, 9, 22].

In light of these contrasting results, we directly investigated in this work the physical properties of the PM lipid bilayer, comparing infected and non-infected cells. To this aim, we used avian and human IAV strains, as well as two cell lines. The previously observed increase of the transport of cholesterol-/sphingolipid-rich vesicles [48] and GPL-conjugated LC3-positive vesicles [37, 41-43] to the PM and their subsequent fusion with the PM might have an impact on membrane organization, composition, fluidity and membrane protein dynamics. For this reason, we used a fluorescence membrane charge sensor (MCS) to monitor changes in the electrostatic potential at the inner leaflet of the membrane [49]. Second, we quantified the influence of IAV infection on PM fluidity by using solvatochromic dyes (Laurdan and Di-4ANEPPDHQ) which are influenced by lipid packing, membrane hydration and lipid composition [50, 51]. Finally, we applied scanning fluorescence spectroscopy (sFCS) to monitor the dynamics of different membrane-associated proteins [52-55]. Our findings indicate that infection by either IAV strains might modulate the lipid composition of the PM and lipid-lipid interactions in both cell models. These changes in membrane properties have a direct effect also on membrane protein dynamics.

## 2. Materials and Methods

### 2.1. Plasmids

The plasmid for FRET analysis, MCS+ [49], was acquired from Addgene (gift from Katharina Gaus, Addgene plasmid #90412). All plasmids for sFCS analysis encoded the monomeric enhanced green fluorescent protein (mEGFP) which was fused to the C-terminus of a myristoylated and palmitoylated peptide (mp-mEGFP), avian influenza A/FPV/Rostock/1934 virus hemagglutinin (HA-mEGFP, Addgene plasmid #127810), or to the N-terminus of a glycosylphosphatidylinositol (GPI)-anchor (GPI-mEGFP, Addgene plasmid #182866) and were previously described [42, 56]. A schematic overview of the localization of each construct within the PM is provided Figure S1A and F.

### 2.2. Cell culture, transfection and infection

Madin-Darby canine kidney type II (MDCK II) cells (ECACC 00062107, European Collection of Authenticated Cell Cultures, Porton Down, UK), chicken embryonic fibroblast cell line DF1 (ATCC number: CRL–12203, kindly provided by Andreas Herrmann, Humboldt University Berlin, Germany) and human embryonic kidney (HEK) cells from the 293T line (CRL-3216TM, purchased from ATCC, Kielpin Lomianki, Poland) were maintained in phenol red-free, high glucose Dulbecco’s modified Eagle’s medium (DMEM) supplemented with 2 mM L-glutamine, 100 U/mL penicillin, 100 µg/mL streptomycin and 10% fetal bovine serum in a humidified incubator at 37°C and 5% CO_2_ atmosphere. Cells were passaged every 2–4 days, until passage 15. All cell culture products were purchased from PAN-Biotech (Aidenbach, Germany).

For imaging experiments, 35-mm dishes (CellVis, Mountain View, CA, USA) with an optical glass bottom (#1.5 glass, 0.16–0.19 mm) were coated with 0.01% (w/v) poly-L-lysine (molecular weight [MW] 150,000–300,000 Da, Sigma-Aldrich, Munich, Germany) for four hours at 37°C and rinsed three times with Dulbecco’s phosphate-buffered saline containing Mg^2+^/Ca^2+^ (DPBS_+/+_; PAN-Biotech, Aidenbach, Germany) before cell seeding. Cells were seeded 24 h prior to transfection and infection at a density of 6 × 10^5^ cells per dish.

For FRET and sFCS measurements, cells were transfected four hours prior infection with Turbofect® according to the manufacturer’s protocol (Thermo Fisher Scientific, Waltham, MA, USA) by using 100 ng pDNA per dish for the membrane-associated proteins (mp-mEGFP and GPI-mEGFP) and the FRET-sensor (MCS+) or 600 ng pDNA per dish for the transmembrane glycoprotein, HA-mEGFP. Briefly, pDNA was pre-incubated with 2 µL of reagent in a final volume of 50 µL serum-free medium and was added dropwise to cells after incubation for 20 min at room temperature.

Before single-round infections with the avian influenza A/FPV/Rostock/1934 virus mutant 1 (FPV, H7N1, kind gift from Michael Veit, Free University Berlin [57]) and the human influenza A/WSN/1933 virus (WSN, H1N1, kind gift from Andreas Herrmann, Humboldt Universität zu Berlin, Germany), cells were washed three times with DPBS_+/+_ and then infected with a multiplicity of infection (MOI) 5 in DMEM containing 0.2 % (w/v) Bovine Serum Albumin (BSA; Sigma Aldrich, Taufkirchen, Germany), 2 mM L-glutamine, 100 U/mL penicillin, and 100 µg/mL streptomycin. Cells were first incubated for 15 min on ice and then for 45 min in a humidified incubator at 37°C and 5% CO_2_ atmosphere. Afterwards, cells were rinsed three times with DPBS_+/+_ and fresh infection medium was added. Cells were further maintained under standard growth conditions until the measurements (∼ 16 hours post infection (hpi)). This time point has been chosen in agreement with previous studies (monitoring infection after 12, 18 and 24 h [7]) and also with our results showing that M1 is efficiently recruited to the PM after ca. 16 hours [46]. Virus propagation and titration were performed in MDCK II cells as previously described [46].

### 2.3. Alteration of PM properties to obtain control samples

For FRET measurements, cells were treated with lipid vesicles containing the anionic phospholipid 1,2-dioleoyl-sn-glycero-3-phospho-L-serine (DOPS, purchased from Avanti Polar Lipids, Inc., Alabaster, AL, USA) to increase the concentration of negative-charged lipids at the PM and, thus, function as positive control [49]. Lipids were dissolved in chloroform at the concentration of 1 mM, dried on the walls of a glass vials under nitrogen gas and stored at -20°C until use. Prior to each experiment, the lipid film was rehydrated with DPBS without Ca^2+^/Mg^2+^ (DPBS_-/-_) to a final concentration of 0.1 mM and vigorously vortexed until multilamellar vesicles (MLV) were formed. The MLV suspension was then sonicated to clarity in a bath sonicator to form small unilamellar vesicles (SUVs). DOPS vesicles were added to the cells after three washing steps with DPBS_-/-_ and incubated for 30 min at room temperature. Afterwards, cells were washed with DPBS_-/-_ and fresh culture medium was added before starting the measurement.

For GP measurements, cells were treated with 10 mM methyl-beta cyclodextrin (MbCD; Sigma-Aldrich, Darmstadt, Germany) in serum-free DMEM after three washing steps with DPBS_+/+_ to obtain a control sample featuring an increase in membrane fluidity, consequent to cholesterol depletion. After incubation for four hours in a humidified incubator at 37°C and 5% CO_2_ atmosphere, cells were washed three times with DPBS_+/+_ before labelling with a fluorophore.

### 2.4. Membrane labelling with Laurdan and Di-4-ANEPPDHQ

The membrane probes Di-4-ANEPPDHQ and Laurdan (6-dodecanyl-2-dimethylaminonaphtalene) were obtained from Molecular Probes (Eugene, OR, USA) and dissolved in ethanol and dimethyl sulfoxide (DMSO, Sigma-Aldrich, Darmstadt, Germany) at the desired concentrations, respectively. Aliquots of 2 mM stock solutions were stored at -20°C until use. Before each experiment, membrane probes were diluted in serum-free and phenol red-free DMEM to a final concentration of 1 µM for Di-4-ANEPPDHQ and 5 µM for Laurdan (the final concentration of DMSO was kept below 0.5% v/v). The probes were added to the cells after three washing steps with DPBS_+/+_ and incubated for 10 min at room temperature. Afterwards, cells were washed twice with DPBS_+/+_ and fresh serum-free and phenol red-free DMEM were added to the cells before imaging.

### 2.5. Confocal spectral imaging

Spectral imaging was performed on a Zeiss LSM 780 system (Carl Zeiss Microscopy GmbH, Oberkochen, Germany) equipped with a Plan-Apochromat 40×/1.2 Korr DIC M27 water immersion objective and a 32-channel gallium arsenide phosphide (GaAsP) detector array. The excitation sources were a 405 nm diode laser (for Laurdan) and a 488 nm Argon laser (for Di-4-ANEPPDQH and the FRET-sensor). Images were collected with detection ranges between for 419 and 610 nm for Laurdan and 499 and 695 nm for Di-4-ANEPPDQH and FRET-sensor in 8.9 nm increments after passing 405/625 nm or 488 nm dichroic mirrors, respectively. Laser power was adjusted so that no pixel saturation occurred. For image acquisition, ten frames were taken with a frame resolution of 128 × 128 pixels, a pixel dwell time of 50.4 µs and a pinhole size of 1 AU to reduce out-of-focus fluorescence. All measurements were performed at room temperature (22 ± 1°C) and images were acquired at the equatorial plane of the cell.

### 2.6. FRET analysis

Confocal spectral images were analyzed as previously described [42, 49, 58]. To quantify the FRET signal, several regions of interest (ROIs) were manually defined at the PM of several cells and emission spectra were computed from the mean normalized fluorescence intensities over all pixels of each ROI, after the application of an intensity threshold (set as 1/5 of the maximum intensity over all wavelengths). Additionally, the normalized pixel intensity values within the ROIs were used to calculate the red-green intensity ratio (RG ratio). The red spectral range was set 601-619 nm and the green from 512-539 nm. Hence, the RG ratio is defined as:

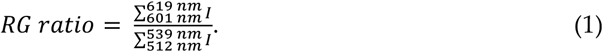

RG ratio values were then either plotted as RG ratio maps or used for the normalized occurrence histograms of all selected pixels. The RG ratio distributions for these samples range from 0 to ca. 1, corresponding to low to high FRET levels. A schematic overview of the FRET analysis is provided in Figure S1 A-B.

All calculations were performed using a custom-written MATLAB code (The MathWorks, Natick, MA, USA). The mean normalized intensity spectra were visualized by using GraphPad Prism vs. 9.0.0 (GraphPad Software, LCC, San Diego, CA).

### 2.7. GP index analysis

Spectral GP measurements were analyzed as previously described [42, 54, 58]. Briefly, an average fluorescence intensity image over 10 frames was calculated before defining ROIs at the PM. After the application of an intensity threshold (set as 1/5 of the maximum intensity over all wavelengths), pixel intensities of each channel were used to calculate the average fluorescence intensity spectrum over all pixels within this mask. Moreover, the generalized polarization (GP) index was calculated from a pixel-wise analysis and the obtained values were used to obtain the GP map, the normalized occurrence histograms of all selected pixels and to compute the average GP value over the entire ROI. For the calculation of the GP values, the wavelengths relative to ordered and disordered phases were previously identified for each probe, by measuring standard giant unilamellar vesicles (GUVs, Figure S1E). In this study, spectral ranges of 420-450 nm (with enhanced emission from ordered bilayers) and 520-560 nm (showing enhanced emission for disordered bilayers) were set for Laurdan. 545-585 nm and 625-690 nm were chosen for Di-4-ANEPPDQH. The sum of the normalized intensity values for each range (I_o_ and I_d_, respectively) was then used for the GP index calculation as followed:

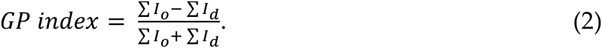

The GP index ranges from -1.0 (very fluid) to 1.0 (gel-like). A schematic overview of the GP index analysis is provided in Figure S1C-E.

All measurements were analyzed with custom-written MATLAB code (The MathWorks, Natick, MA, USA). The mean normalized intensity spectra were visualized by using GraphPad Prism vs. 9.0.0 (GraphPad Software, LCC, San Diego, CA).

### 2.8. sFCS analysis

Scanning FCS experiments were performed on a Zeiss LSM780 system (Carl Zeiss Microscopy GmbH, Oberkochen, Germany) as previously described [54, 56, 59]. Briefly, the samples were excited with 488 nm Argon laser (≈3 µW) through a Plan-Apochromat 40×/1.2 Korr DIC M27 water immersion objective. The fluorescence signal was collected between 499 and 600 nm with a 32-channel GaAsP detector array after passing through a 488 nm dichroic mirror. The pinhole size was restricted to an airy unit of one to minimize out-of-focus signal. Line scans of 256 × 1 pixels and pixel size ≈ 80 nm were acquired perpendicular to the PM with a scan time of 472.73 µs. Typically, 400,000 lines were acquired in photon counting mode. All measurements were performed at 22 ± 1°C.

Prior to each experiment, the confocal volume was calibrated by performing a series of point FCS measurements with a 30 nM Alexa Fluor® 488 solution (AF488, Thermo Fischer, Waltham, MA, USA) at the same excitation power and beam path used for sFCS measurements. For that, the fluorescence signal was optimized at first by adjusting the collar ring of the objective and the pinhole position to the maximal count rate. Then, FCS measurements (15 repetitions of 10 s) at five different positions were performed and the data was fitted using a 3D diffusion model including a triplet contribution in order to calculate the structure parameter S (ratio of the vertical and lateral dimension of the confocal volume, typically between 6 and 8) and the diffusion time τ_*d*_(usually, ≈ 30 ± 2 µs). The measured average diffusion time and a previously determined diffusion coefficient (*D*_488_ =435µ*m*^2^*s*^−1^) [60] were used to calculate the waist ω_0_ of the confocal volume (usually, ≈ 230 ± 10 nm).

Scanning FCS data were exported as TIFF files, imported and analyzed in MATLAB (The MathWorks, Natick, MA, USA) using a custom code as previously described [46, 54, 56]. Briefly, all scanning lines were aligned and divided into blocks in which the lines were fitted with a Gaussian function in order to define the membrane position. Subsequently, pixels of each line were integrated to provide the membrane fluorescence time series *F(t)*. Photobleaching was corrected by a two-component exponential fit function [61]. Afterwards, the normalized autocorrelation function (ACF) was calculated using the following Equations (3) and (4):

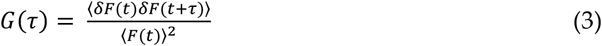

where

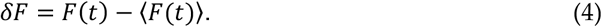

The ACF was calculated segment wise (each ca. 20 s long) and segments with extreme alterations were removed before averaging the ACFs. Finally, a two-dimensional diffusion fitting model and the structure parameter *S* from the calibration were used to analyze the ACF [62], as described in Equation (5):

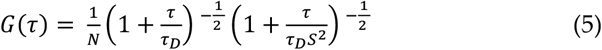

where τ_*D*_ represents the diffusion time and N the number of particles. The diffusion coefficient *D* was then calculated using the waist ω_0_ from the calibration as follows:

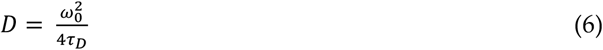

A schematic overview of the sFCS analysis is provided in Figure S1G.

### 2.9. Statistical analysis

Data from at least two independent experiments were pooled, analyzed and visualized using a self-written R script (R Foundation for Statistical Computing, Vienna, Austria) built from common packages (*rstatix, fBasics, ggplot2* and *ggpubr*). Data is displayed as box plots with single data points corresponding to measurements in single cells. Median values and whiskers ranging from minimum to maximum values are displayed. The corresponding descriptive statistics for each plot are summarized in supplementary tables (see supplementary material). The *p* values are provided in each graph and figure captions. Statistical significance was tested by using D’Agostino-Pearson normality test followed by one-way ANOVA analysis and Tukey’s multiple comparisons test.

## 3. Results

### 3.1. Infection increases the negative surface charge of the inner leaflet of the PM

Previous studies have shown that the viral proteins HA and M1 can modulate the clustering of PIP2 in the PM [31, 32]. Moreover, it was reported that the apical transport of M2 is carried out via PS-conjugated LC3-positive vesicles [41-43, 63]. Both observations could have an impact on the membrane composition and, specifically, the amount and lateral organization of anionic lipids.

First, we investigated to what extent the infection induced apoptosis which, in turn, could lead to PS flipping to the outer leaflet of the membrane, as it was reported for late infection states [64, 65]. Trans-bilayer rearrangement of PS might in fact influence the quantification of membrane charge distribution [66, 67]. We have therefore characterized the state of the cells using PI and Annexin V to determine cell viability and apoptosis-induced PS flipping to the outer leaflet of the PM [64, 67]. Infection status and total cell numbers were determined via immunofluorescence and Hoechst 33342 staining, respectively (Figure S2, Table S1). We used H_2_O_2_/Saponine-treated cells as positive control for apoptosis and cell death [68, 69]. The FPV infection efficiencies in HEK293T and DF1 samples were 81.9 ± 4.9 % and 64.7 ± 8.3 % (mean ± SD). The values for WSN were 90.6 ± 3.6 % and 83.1 ± 14.7 % (mean ± SD) (Figure S2A-B, E). We observed no significant induction of apoptosis or PS translocation 16 h after infection with FPV or WSN, in either cell line (Figure S2A-D).

Next, we used a fluorescence membrane charge sensor (MCS+) to monitor changes of the electrostatic potential at the inner leaflet of the membrane [49] in non-infected (MOCK), FPV-/WSN-infected and DOPS-SUV-treated HEK293T and DF1 cells (Figure 1). FRET measurements to quantify the membrane potential were carried-out via spectral imaging, instead of the standard, filter-based method [49]. The spectral imaging approach and detection in the photon-counting mode were previously shown to provide more information and to improve the accuracy and sensitivity of the measurements [70, 71]. For the FRET analysis, ROIs at the equatorial plane of the cells were chosen and the RG ratio quantified. High RG ratio values correspond to a relative higher fluorescence emission peak in the longer-wavelength region (i.e., more efficient FRET between the MCS+ domains) and, therefore, a higher negative membrane potential at the inner leaflet of the membrane. We used DOPS-treated cells as a positive control, since it was shown that DOPS is quickly internalized and transported to the inner leaflet of the PM with a half time of a few minutes [72, 73]. The sensor MCS+ was expressed with similar efficiency in all cell lines and differently treated samples (data not shown). For both cell lines, we observed an increase of the second emission maximum upon virus infection or DOPS-SUV-treatment (Figure 1A), which resulted in ∼2-fold higher RG ratio values (Figure 1B-C and S3, Table S2). Interestingly, no significant differences between IAV strains or cell types were detectable. In summary, these findings indicate a significant increase of the available negative charge at the inner leaflet of the PM in FPV- and WSN-infected cells, independent of their host cell type.

**Figure 1:**
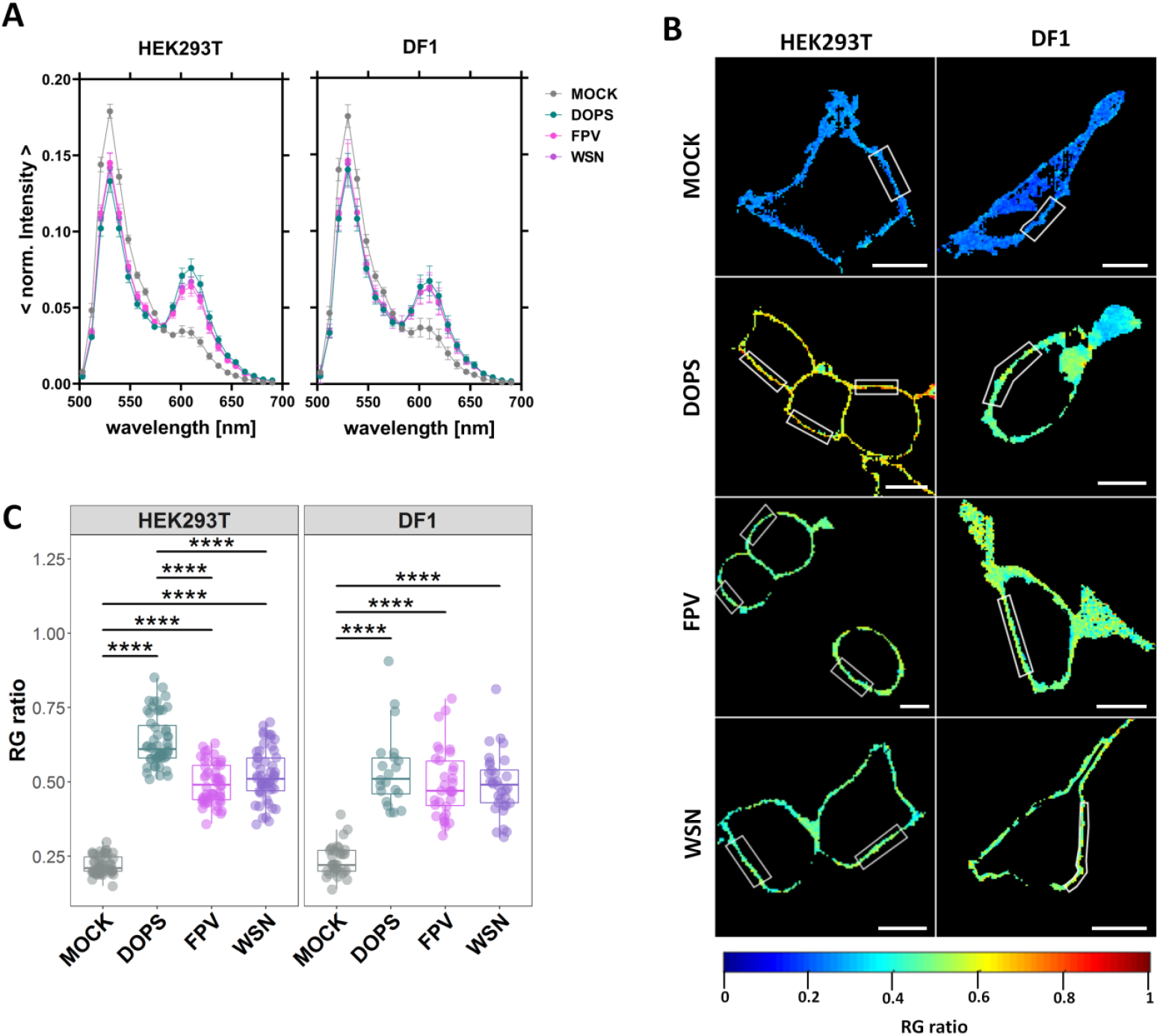
Increase of negative surface charge at the inner leaflet of the PM in infected cells. HEK293T and DF1 cells were either: non-infected (MOCK), treated with DOPS-SUV (DOPS, positive control), infected with FPV or with WSN influenza A strains. All cells were expressing the FRET-sensor MCS+ and emission spectrum images (22 spectral channels from 499 nm to 695 nm) were acquired 16 hpi using 488 nm excitation. **(A)** Average normalized emission spectra of all the selected regions of interest (ROI) at the equatorial plane of HEK293T and DF1 cells expressing MCS+, following the indicated treatment. Data are represented as mean ± SD from 50-55 HEK293T cells and 21-33 DF1 cells. **(B)** Representative ratiometric FRET images (RG ratio, pseudo-colored as indicated by the color scale) of HEK293T and DF1 cells expressing MCS+. White rectangles represent examples of ROIs at the PM selected for FRET quantification. Scale bars represent 10 µm. **(C)** RG ratio derived from the average intensity spectra of each cell type for the indicated treatment. Data from two separate experiments were pooled, plotted and analyzed using one-way ANOVA Tukey’
ss multiple comparison test (**** p < 0.0001). Each data point represents the average value measured for a ROI at the PM in one cell (Table S2).

### 3.2. IAV infection increases lipid packing in the plasma membrane lipid bilayer

Multiple studies have shown that the membrane of influenza A virions contains high levels of sphingolipids and cholesterol, a modified GPL composition and is more ordered compared to the host cell membrane [5, 7, 19, 21, 22, 74, 75]. Little is known about alterations in the physical properties (e.g., order, lipid packing) of the host cell membrane. Therefore, we investigated the effect of IAV infection on membrane order directly in the PM of living cells, using solvatochromic probes (i.e., Laurdan and Di-4ANEPPDHQ). The spectroscopic properties of these dyes depend on the local membrane environment [50, 51]. Specifically, their emission spectra exhibit a blue-shift when they localize in an ordered, more apolar environment, such as a liquid-ordered phase or a “lipid-raft” domain [50, 51]. This shift can be quantified by calculating the GP value, which involves a ratiometric analysis of the fluorescence intensity in two spectral regions [58]. Higher GP values indicate a higher fluorescence intensity in the shorter-wavelength region, which corresponds to a higher degree of membrane order and lipid packing [51].

We used GUVs with varying lipid compositions as reference for the behavior of each dye in solid ordered gel (L_ß_) phase and liquid-disordered (L_d_) phase membranes. With such reference samples, we could reliably test our experimental conditions, the GP analysis pipeline, and the phasor approach (Figure S1E and S4). DLPC GUVs (consisting of a bilayer in the L_d_ phase) were significantly distinguishable from DPPC GUVs (bilayer in the L_ß_ phase), using either Laurdan or Di-4-ANEPPDHQ. In agreement with previous reports, the spectral shift of Di-4-ANEPPDHQ is less dramatic than that observed for Laurdan [50, 58, 76-78].

The investigation of changes in lipid packing was carried out in non-infected cells (MOCK), FPV- and WSN-infected cells and MbCD-treated cells (Figure 2, S5-S6, Table S3-S4). MbCD-treated cells were used as control, since it is known that cholesterol depletion reduces membrane order [58, 67]. In all cases, PM regions at the equatorial plane of the cells were chosen and the GP index was quantified from the spectral information of each pixel. We observed in both cell lines a shift in fluorescence emission towards the shorter-wavelength region upon FPV- and WSN-infection (Figure 2A, D). This shift was less pronounced in Di-4-ANEPPDHQ-stained cells, in agreement with control experiments in GUVs (Figure S4).

**Figure 2:**
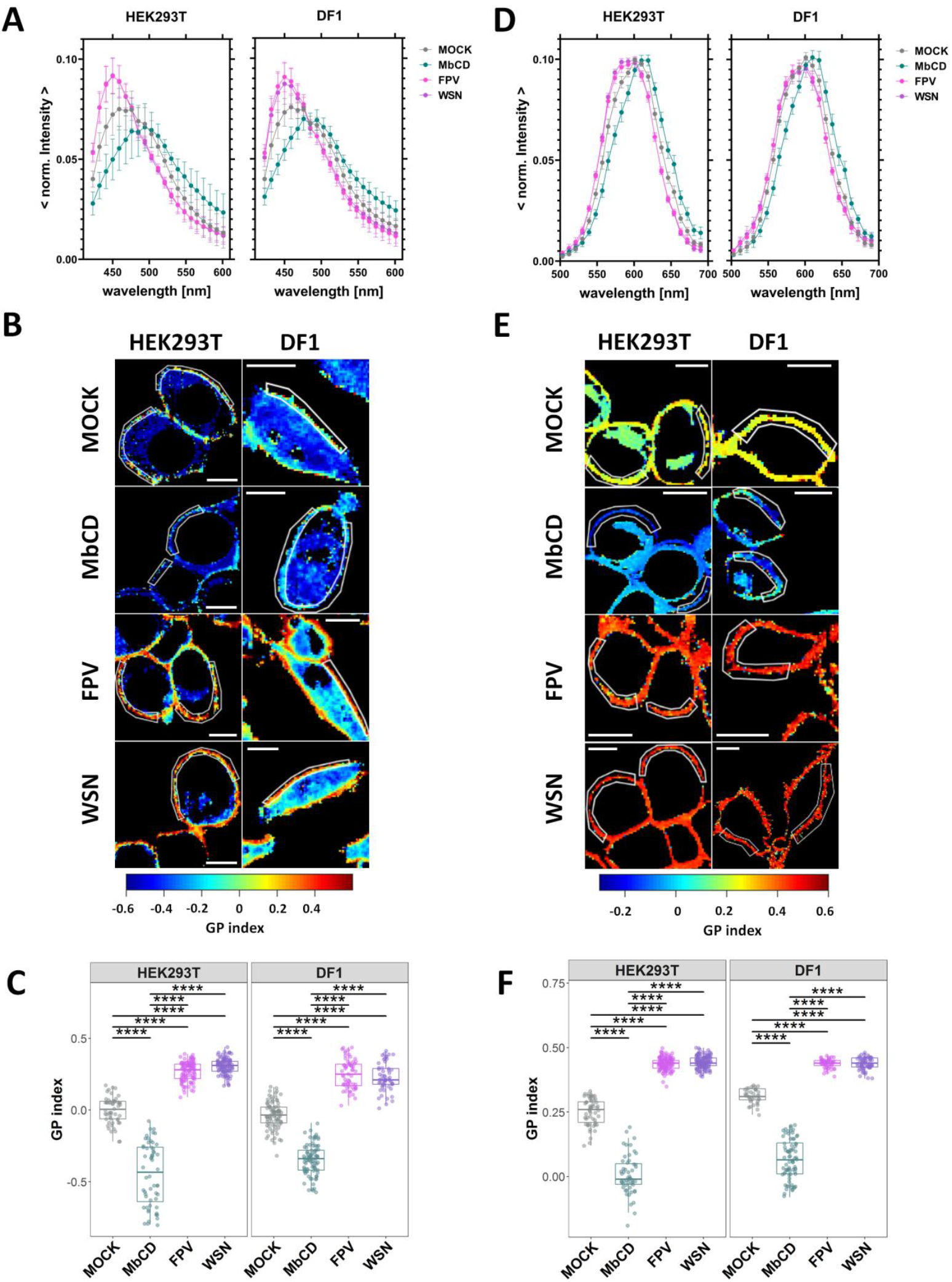
Increase of lipid packing of the PM upon IAV infection. HEK293T and DF1 cells were either: non-infected (MOCK), treated with methyl-β-cyclodextrin (MbCD), infected with FPV or with WSN influenza A strains. All cells were labelled with the solvatochromic probes Laurdan **(A-C)** and Di-4-ANEPPDHQ **(D-F)**, and then imaged 16 hpi. Averaged normalized fluorescence emission spectra of all selected regions of interest (ROI) at the equatorial plane of HEK293T and DF1 cells stained with Laurdan **(A)** or Di-4-ANEPPDHQ **(D)**, for the indicated treatment. Data are represented as mean ± SD of 52-110 cells stained with Laurdan and 36-127 cells stained with Di-4-ANEPPDHQ (Table S3 and S4). Representative ratiometric GP images (GP index, pseudo-colored as indicated by the color scale) of HEK293T and DF1 cells stained with Laurdan **(B)** or Di-4-ANEPPDHQ **(E)**. White lines represent examples of ROIs at the PM selected for GP index quantification. Scale bars represent 10 µm. GP index derived from the average intensity spectra from Laurdan-**(C)** or Di-4-ANEPPDHQ-stained **(F)** cells for each cell type and indicated treatment. Data from three separate experiments were pooled, plotted and analyzed using one-way ANOVA Tukey’
ss multiple comparison test (**** p < 0.0001). Each data point represents the average value measured for a ROI at the PM in one cell (Tables S3 and S4).

MbCD-treated cells showed as expected, for all cell types, a red-shifted spectrum [58, 67]. Moreover, the alternative representation via phasor plots showed a clear “clockwise” shift (see also Figure S4C) for infected cells (Figure S5), confirming stronger lipid-lipid interactions in these samples. Finally, our data indicate that the observed PM ordering effect consequent to IAV infection does not depend on IAV strain, cell type (Figure 2C and F, S5-S6, Table S3 and S4), or degree of labelling (data not shown).

### 3.3. IAV infection reduces membrane protein dynamics

Next, we aimed to determine whether the changes induced by IAV infection in the context of PM physical properties (see previous paragraphs) and lipid composition [6, 7, 9, 22] have an effect on the diffusive dynamics of trans-membrane proteins. Therefore, we quantified the in-plane diffusion of three membrane(-associated) proteins: i) a model for a protein associated to the inner leaflet of the PM (mp-mEGFP), ii) a model for a protein associated to the outer leaflet of the PM (GPI-mEGFP) and iii) a model of a trans-membrane protein (HA-mEGFP). Measurements were carried out in non-infected and FPV-/WSN-infected (16 hpi) HEK293T and DF1 cells. Quantification of the diffusion dynamics of the fluorescently labelled proteins was performed via sFCS measurements perpendicular to the membrane, at the equatorial plane of the cells. Representative cell images and ACFs obtained in HEK293T cells are shown in Figures S7 and S8, respectively.

Quantitative analysis of the ACFs indicated that both membrane-associated proteins diffuse in the PM of non-infected cells with a diffusion coefficient (D) of ≈ 1.1 µm^2^/s, while the transmembrane protein exhibit slower dynamics (D ≈ 0.4 µm^2^/s, Figure 3, Table S5). Both results are in line with previous experiments [46, 56, 79]. In FPV- and WSN-infected cells, we observed a decrease of diffusive dynamics for both mp-mEGFP (ca. 2-fold change) and GPI-mEGFP (ca. 1.8-fold change), for both cell lines. A decrease of mobility for the model transmembrane protein, HA-mEGFP, was also observed, albeit smaller (ca. 1.4-fold) and only in avian DF1 cells infected with either IAV strain.

**Figure 3:**
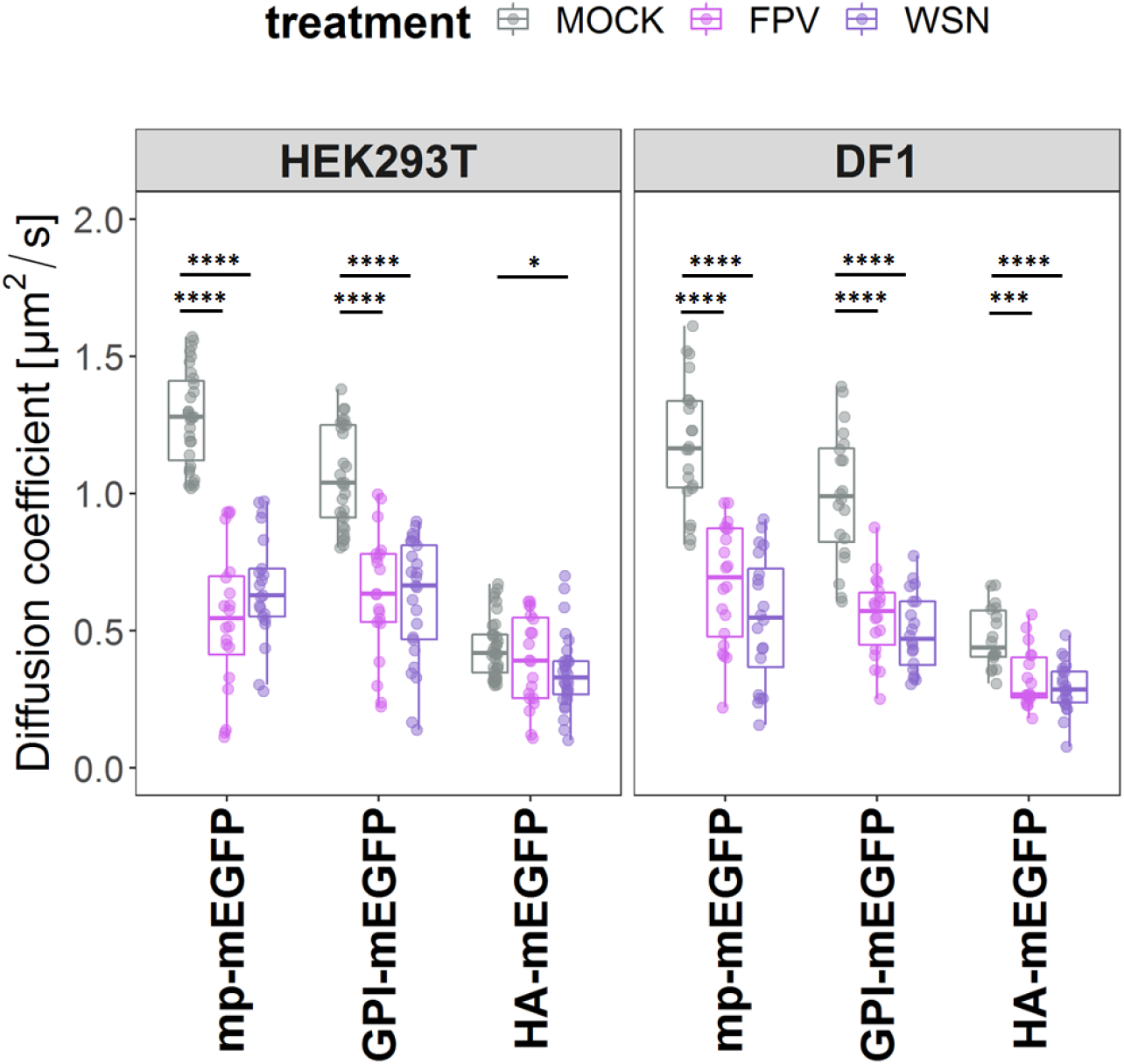
Decrease of membrane protein diffusion upon IAV-infection. Quantitative analysis of protein diffusion via fluorescence correlation spectroscopy (sFCS) in non-infected (MOCK) and FPV-/WSN-infected HEK293T and DF1 cells expressing three model proteins labelled with green fluorescent proteins (mEGFPs) and associated to the plasma membrane (PM). Specifically, we investigated: i) a construct anchored to the inner-leaflet of the PM via a myristoylated and palmitoylated (mp) peptide (mp-mEGFP), ii) a construct anchored to the outer-leaflet of the PM via a glycosylphosphatidylinositol (GPI) anchor (GPI-mEGFP) and iii) one representative transmembrane protein, i.e. the influenza envelope protein hemagglutinin (HA-mEGFP). Measurements were performed 16 hpi. The box plots show the diffusion coefficients calculated from sFCS diffusion times. Data from three separate experiments were plotted and analyzed using one-way ANOVA Tukey’
ss multiple comparison test (* p < 0.05, *** p < 0.001, **** p < 0.0001). Each data point represents the value measured at the PM in one cell (Table S5).

## 4. Discussion

Previous studies have shown that IAV infection induces changes in the lipid metabolism of infected cells [6, 7, 9]. Therefore, the concentration of specific lipids in cellular membranes might change as a consequence of infection [6, 7, 9]. For example, an IAV-induced increase in saturated lipids [5-7], cholesterol/sphingolipids [6, 7, 9] or anionic lipids [7, 9] might significantly alter the physical properties of the PM, including local order of the lipid bilayer and its surface charge at the inner leaflet side. Such alterations, in turn, might affect protein-protein and protein-lipid interactions [80]. Here, we have quantified the effect of IAV infection on the properties of the PM and, specifically, its fluidity, structural order, and surface charge.

One finding of this study is the enhancement of the negative surface electrostatic potential at the inner leaflet of the PM, upon IAV infection, as demonstrated using a FRET-based membrane charge sensor. The simplest explanation for this result is an increase of the local concentration of anionic lipids in the inner leaflet. This idea is supported by the fact that the concentration of, e.g., PS in the membranes of infected cells or in the viral envelope was shown to be higher than in non-infected cells [5, 7, 9, 21]. On the other hand, it must be kept in mind that our results might also be compatible with the electrostatic potential being altered by, e.g., changes in cytoplasm ionic strength/composition or, more likely, alterations in lipid/protein lateral organization [31, 44]. In the latter case, it is reasonable that clustering of negative lipids or unbinding of positively charged proteins interacting with anionic lipids might alter the (local) effective electrostatic potential. In any case, such an increase in the amount or “availability” of anionic lipids at the inner leaflet might indeed relate to the organization of several viral components. For instance, it was shown that M1 is recruited by M2 to the PM and that M1 co-clusters with PS as well as with PIP2 [32, 44, 46]. Similar observations were made in the case of HA and both proteins contain polybasic residues which play a role in their interaction with anionic lipids and membrane localization [31, 44, 45]. Additionally, viral protein transport occurs via LC3, a protein that interacts with anionic lipids, especially in IAV-infected cells [41-43]. Finally, also genome packing is regulated by anionic lipids at the PM (i.e., PIP2) interacting with viral RNA and the IAV nucleoprotein NP [81].

Overall, these observations suggest that alterations in the local concentration of anionic lipid at the inner leaflet or, more in general, alterations of the negative electrostatic potential of infected cells might modulate viral assembly and release. Our results show for the first time that such alterations are indeed directly observable at the PM of infected cells, independent of IAV strain or cell type. Of note, this might be a general phenomenon, common also to other viral infections. For example, reduction in PS and PIP2 levels in mammalian cells hinders the recruitment of VP40 protein to the PM and its oligomerization, thereby inhibiting Ebola virus/Marburg virus assembly and egress (reviewed in [82]). Rearrangements of lipids at the PM, such as PIP2 and cholesterol, were also observed during the assembly of the human immunodeficiency virus (HIV) Gag protein (reviewed in [82]).

Another finding reported in this work is the enhancement of lipid-lipid interactions in the PM of infected cells. Specifically, we have used Laurdan and Di-4-ANEPPDHQ to quantify the impact of IAV infection on membrane fluidity and lipid packing. These two fluorophores are commonly used to probe membrane order [50, 51, 58, 78] and are influenced, each in a specific way, by several factors including: cholesterol content (in connection to glycerol backbone dynamics), membrane hydration (in connection to lipid internal motions and hydrogen bond network dynamics) [77, 83] and lipid phase behavior [50, 78]. Furthermore, it was reported that while Laurdan is a reliable probe for membrane order, Di-4-ANEPPDHQ is influenced by cholesterol and membrane potential [50]. Interestingly, our results indicate that both probes report similar alterations in the PM of IAV-infected cells. Taking into account also previous lipidomic analyses [6, 7, 9], the simplest explanation for these data is indeed an increase in the (local) concentration of cholesterol and/or saturated lipids (including e.g., sphingolipids). Of interest, this interpretation is compatible with our results regarding the possible increase of negatively charged lipids: on one hand, an increase in cholesterol can be accompanied by higher levels of the anionic lipid PIP2 in cellular membranes (reviewed in [84]); on the other hand, a recent report linked the presence of PS to enhanced interleaflet coupling in model membranes and, as a consequence, an increase in membrane stiffness [85]. Furthermore, an increase in PM cholesterol concentration might, at least, partially explain the enhanced M2 clustering (see Figure S2, and compared to transfected cells [46]) which is, in fact, modulated by cholesterol [35, 39, 40]. This issue is the object of current studies in our laboratory.

As a consequence of the altered lipid composition and increased structural order, the diffusive dynamics of membrane components might be hindered. The in-plane mobility of transmembrane proteins is tightly connected to their function [86] and affects several cellular processes [87]. To study membrane dynamics in infected cells, we used two model fluorescent proteins associating with the inner leaflet of the PM (mp-GFP) or to its outer leaflet (GPI-GFP). Both markers diffused significantly slower after infection, in agreement with the increased membrane order that was detected upon infection, via e.g. Laurdan-based assays. This effect was even stronger at later infection time points (data not shown). The decreased fluidity of both leaflets might be due to alterations of either local lipid composition (i.e., restricted to one specific leaflet and transmitted to the other leaflet through enhanced interleaflet coupling [85, 88]) or general bilayer composition (e.g. cholesterol concentration, affecting both leaflets [88]). Interestingly, the effect of infection on the diffusion dynamics of HA was less pronounced than for the other test proteins. This could be due to the fact that the diffusive dynamics of transmembrane proteins are determined also by other factors, such as interaction with the cytoskeleton, rather than just lipid bilayer viscosity [89-91]. Moreover, in the case of HA, it might be possible that this protein is enriched in specific domains [27, 30, 92] with a local lipid composition which is not significantly altered during infection. Nevertheless, our findings indicate that the diffusive dynamics of membrane-associated proteins are hindered in general by the decrease in membrane fluidity and/or increase in lipid packing. In general, such alterations in the membrane order parameters might originate from an overall re-organization of the membrane components or might be caused by the presence of, e.g., locally-ordered membrane domains [11] from which the virus can efficiently assemble and bud.

Although our study focuses on the consequences of the increased lipid-lipid interactions in the PM of infected cells, it is interesting to speculate about the possible mechanisms leading to, e.g., alterations in membrane compositions during IAV infection. So far, the viral proteins NA and M2 have been reported to directly alter the fatty acid metabolism of host cells [93]. Also, it appears that HA and NA are transported to the PM via cholesterol-/sphingomyelin-rich vesicles that might alter the composition of the target membrane [13, 15, 18, 20, 21, 25]. Of interest, enrichment of cholesterol and saturated ordered-inducing lipids at the PM (or, specifically, at the budding site) might be important for the environmental stability of the virus and virus morphology [23, 24, 94, 95].

## 5. Conclusions

In this study, we provide evidence for IAV-induced alterations of PM dynamics and structural organization in infected cells. To the best of our knowledge, we demonstrate for the first time that IAV infection induces a decrease in membrane fluidity, an increase in lipid packing and an enhancement of the negative electrostatic potential at the PM inner leaflet (probably caused by increased local concentrations of anionic lipids). Moreover, our study highlights the potential of combined biophysical methods to investigate membrane properties at the single cell level, from multiple points of view, with the aim of better understanding virus-host interactions. These techniques can also be utilized in future studies to explore the effects of specific agents targeting lipid metabolism and host cell PM properties on virus egress and replication. Additionally, it might be possible to shed light on the role of specific viral proteins that influence membrane physical properties during virus assembly.

## Supplementary Materials

Supporting Materials and Methods, Table S1: Overview of the Annexin V, cell viability and infection status analysis for HEK293T and DF1 cells, Table S2: Overview of the RG ratio from the FRET analysis for HEK293T and DF1 cells, Table S3: Overview of the GP index analysis with Laurdan for HEK293T and DF1 cells, Table S4: Overview of the GP index analysis with Di-4-ANEPPDHQ for HEK293T and DF1 cells, Table S5: Overview of the diffusion coefficient values [µm2/s] from the sFCS analysis for HEK293T and DF1 cells, Figure S1: Schematic overview of the experimental setup for the analysis of the enrichment of negatively charged lipids at the inner leaflet of the PM via FRET (A-B), changes in membrane fluidity via GP index (C-E), and diffusion time of PM-associated peptids/proteins using sFCS (F-G), Figure S2: No induction of apoptosis or PS flip upon FPV or WSN infection in human and avian cells, Figure S3: Quantification of the RG ratio from the FRET measurements at the PM of HEK293T and DF1 cells, Figure S4: GP and spectral phasor analysis of Laurdan and Di-4-ANEPPDHQ labelled GUVs, Figure S5: Spectral phasor analysis of Laurdan and Di-4-ANEPPDHQ labelled HEK293T and DF1 cells, Figure S6: Quantification of the GP index from the membrane fluidity measurements at the PM of HEK293T and DF1 cells with Laurdan and Di-4-ANEPPDHQ, Figure S7: Representative confocal fluorescence images of HEK293T cells expressing mEGFP-tagged proteins, Figure S8: Examples of autocorrelation functions, Data S1: Excel sheet including data for Figure 1A, Figure 1C, Figure 2A,. Figure 2C, Figure 2D, Figure 2F, Figure 3, Figure S2C-E, Figure S4A.

## Author Contributions

Conceptualization, A.P.; methodology, A.P.; software, A.P. and S.C.; validation, A.P.; investigation, A.P.; writing—original draft preparation, A.P.; writing—review and editing, S.C.; visualization, A.P.; supervision, S.C.; project administration, S.C.; Funding Acquisition, S.C. All authors have read and agreed to the published version of the manuscript.

## Funding

This research was funded by the Deutsche Forschungsgemeinschaft (DFG) (grant numbers #254850309 to S.C.).

## Data Availability Statement

The datasets analyzed during the current study are included the Supplementary Materials.

## Code Availability Statement

MATLAB custom-written codes are available from the corresponding author upon reasonable request.

## Acknowledgments

We are grateful to Andreas Herrmann and the members of the Physical Biochemistry group for critical reading of the manuscript.

## Conflicts of Interest

The authors declare no conflict of interest.

## Supporting citations

References **[49, 58, 96-102]** appear in the supplementary material.

## Notes

### Competing Interest Statement

The authors have declared no competing interest.

