## Supplementary Information for "Influenza A virus infection alters lipid packing and surface electrostatic potential of the host plasma membrane"

### Supplementary Material

#### 1. Materials and Methods

*1.1 Cell Surface immunofluorescence and Annexin V/PI staining.*

To assess the infection status (i.e., infected vs. non-infected), cells were washed three times with DPBS$_{+/+}$ 16hpi, before adding the primary antibody, monoclonal mouse anti-influenza A M2, clone 14C2 (#ab5416, Abcam, Cambridge, UK), with a dilution of 1:200 in 0.2 % (w/v) BSA in DMEM. Cells were incubated for one hour in a humidified incubator at 37°C and 5% $CO_2$ atmosphere. After three washing steps with DPBS$_{+/+}$, cells were incubated with 1:1000 diluted secondary antibody (goat anti-mouse AlexaFluor® 647-conjugated; Thermo Fisher Scientific, Waltham, MA, USA) for one hour in a humidified incubator at 37°C and 5% $CO_2$ atmosphere. Cells were subsequently washed three times with DPBS$_{+/+}$ and stained with 7 mM AlexaFluor® 488-labelled Annexin V, 2 µg/mL propidium iodide (PI) and 2 µg/mL Hoechst 33342 for 15 min at room temperature, to assess cellular integrity and the exposure of PS. All probes were purchased from Thermo Fisher Scientific (Waltham, MA, USA). Afterwards, cells were washed once more with DPBS$_{+/+}$ and analysed using a Zeiss LSM 780 system (Carl Zeiss Microscopy GmbH, Oberkochen, Germany). Confocal fluorescence images were acquired with a Plan-Apochromat 40×/1.2 Korr DIC M27 water immersion objective and an image resolution of 512 x 512 pixels. Samples were excited with a 405 nm diode laser (Hoechst 33342), a 488 nm Argon laser (Annexin V), a 561 nm diode laser (PI) and 633 diode laser (M2). Fluorescence was observed between 415 – 502 nm (Hoechst 33342), 490 – 570 nm (Annexin V), 588 – 650 nm (PI) and 650 – 735 nm (M2), after passing a 405/505c dichroic mirror or 488/561/633 dichroic mirror. Finally, all cells were counted and classified via ImageJ (http://imagej.nih.gov/ij/) to calculate the amounts of Annexin V-positive cells (%), cell viability (%) and infection status (%).

Non-infected (MOCK) cells functioned as negative control for the infection status and cells treated with 8 µM $H_2O_2$/0.1% Saponine for 10 min in a humidified incubator at 37°C and 5% $CO_2$ atmosphere, were used as positive control for PS externalization (Annexin V) and cell death (PI).

*1.2 Giant unilamellar vesicles preparation.*

Giant unilamellar vesicles (GUVs) were produced using the electroformation method [1, 2] and were used for microscopic visualization of the GP index and Phasor analysis in $L_o$ and $L_d$ membranes. These measurements served as reference points for the solvatochromic probes described below. We used the following lipids (purchased from Avanti Polar Lipids, Inc., Alabaster, AL, USA): 1,2-dipalmitoyl-sn-glycero-3-phosphocholine (DPPC) as main component of the solid ordered gel ($L_ß$) phase, and 1,2-dilauroyl-sn-glycero-3-phosphocholine (DLPC) as main component of the $L_d$ phase. Briefly, 4 µl of a 2.5 mg/mL lipid solution were homogeneous spread onto two parallel Pt wires mounted in a custom-made cylindrical Teflon chamber and the chloroform was evaporated under a nitrogen stream for five minutes at room temperature. Then, the chamber was filled with a 50 mM sucrose solution in deionized water and the wires were placed in the solution and connected to a voltage generator (AC generator FG 250 D, H-Tronic, Hirschau, Germany). GUV formation was induced by applying a sinusoidal electric field with an amplitude (peak-to-peak) of 2 V and a frequency of 10 Hz for 1 h at room temperature (DLPC) or above the specific transition temperature at 70 °C (DPPC). The fission of GUVs from the wires surface was facilitated by lowering the frequency to 2 Hz and the voltage to 1.3 V for at least 30 minutes. Subsequently, the GUVs were gently mixed 1:1 with 40 mOsm/kg PBS, stained with 4 µM of probes and imaged in a previously coated (0.1 mg/ml bovine serum albumin in 40 mOsm/kg PBS, pH 7.4) 8 well ibidi® glass bottom μ-slide (ibidi GmbH, Gräfelfing, Germany).

*1.3 Spectral phasor analysis*

The fluorescence emission spectra from each pixel of the ROIs within spectral images were transformed into the phasor coordinates ($g(\lambda)$ and $s(\lambda)$) as following:

$$x\ coordinate = g(\lambda) = \frac{\sum_{\lambda} I(\lambda)\cos\left(\frac{2\pi n(\lambda-\lambda_{min})}{\lambda_{max}-\lambda_{min}}\right)}{\sum_{\lambda} I(\lambda)} \quad (1)$$

$$y\ coordinate = s(\lambda) = \frac{\sum_{\lambda} I(\lambda)\sin\left(\frac{2\pi n(\lambda-\lambda_{min})}{\lambda_{max}-\lambda_{min}}\right)}{\sum_{\lambda} I(\lambda)} \quad (2)$$

The coordinates $g(\lambda)$ and $s(\lambda)$ represent the real and imaginary component of the Fourier transformation, respectively. $I(\lambda)$ is the intensity for each wavelength and $n$ is the harmonic number. We restricted our analysis to the first harmonic ($n = 1$) and the conclusions of the analysis remained qualitatively similar setting $n$=2. The $x$- and $y$-coordinates were then plotted in the four-quadrant spectral phasor plot as previously described [3]. The coordinates $g(\lambda)$ and $s(\lambda)$ take values between 1 to -1. The angular position in the phasor plot is proportional to the center of mass and the phasor radius is inversely proportional to the full width at half maximum (FWHM) of the emission spectrum. The advantage of the phasor approach is that small spectral shifts of Laurdan/ Di-4-ANEPPDHQ emission from labelled cells (caused by small changes in the lipid microenvironment) can be more easily resolved. Moreover, the phasor analysis takes into account the entire spectrum (rather than specific wavelengths or intervals) and it is model-free.

All measurements were analyzed with custom-written MATLAB code (The MathWorks, Natick, MA, USA). A schematic overview of the spectral phasor analysis is provided in Figure S1E and representative analysis for GUVs in Figure S4.

**2. Tables**

**Table S1: Overview of the Annexin V, cell viability and infection status analysis for HEK293T and DF1 cells.** Data correspond to Figure S2. n: number of images, iqr: interquartile range, sd: standard deviation, se: standard error of the mean, ci: confidence interval.

| cell type | treatment | n | median | iqr | mean | sd | se | ci |
|---|---|---|---|---|---|---|---|---|
| **% Annexin V** | | | | | | | | |
| HEK293T | MOCK | 16 | 1.1 | 1.9 | 1.4 | 1.6 | 0.4 | 0.8 |
| HEK293T | $H_2O_2$-Saponine | 15 | 86.4 | 10.8 | 86.7 | 7.2 | 1.9 | 4.0 |
| HEK293T | FPV | 15 | 1.7 | 1.5 | 2.2 | 2.0 | 0.5 | 1.1 |
| HEK293T | WSN | 15 | 0.9 | 1.4 | 1.0 | 0.9 | 0.2 | 0.5 |
| DF1 | MOCK | 18 | 3.9 | 6.3 | 3.9 | 3.8 | 0.9 | 1.9 |
| DF1 | $H_2O_2$-Saponine | 15 | 100.0 | 0.0 | 100.0 | 0.0 | 0.0 | 0.0 |
| DF1 | FPV | 15 | 0.0 | 7.5 | 3.8 | 5.2 | 1.3 | 2.9 |
| DF1 | WSN | 15 | 0.0 | 9.6 | 4.5 | 5.1 | 1.3 | 2.8 |
| **% cell viability** | | | | | | | | |
| HEK293T | MOCK | 16 | 99.6 | 1.8 | 98.9 | 1.7 | 0.4 | 0.9 |
| HEK293T | $H_2O_2$-Saponine | 15 | 0.0 | 0.0 | 0.0 | 0.0 | 0.0 | 0.0 |
| HEK293T | FPV | 15 | 100.0 | 0.0 | 99.9 | 0.3 | 0.1 | 0.2 |
| HEK293T | WSN | 15 | 99.1 | 1.4 | 99.0 | 0.9 | 0.2 | 0.5 |
| DF1 | MOCK | 18 | 98.5 | 5.8 | 96.7 | 3.9 | 0.9 | 2.0 |
| DF1 | $H_2O_2$-Saponine | 15 | 0.0 | 0.0 | 0.0 | 0.0 | 0.0 | 0.0 |
| DF1 | FPV | 15 | 100.0 | 0.0 | 98.6 | 3.8 | 1.0 | 2.1 |
| DF1 | WSN | 15 | 100.0 | 9.6 | 94.6 | 6.7 | 1.7 | 3.7 |
| **% infected cells** | | | | | | | | |
| HEK293T | MOCK | 16 | 0.0 | 0.0 | 0.0 | 0.0 | 0.0 | 0.0 |
| HEK293T | $H_2O_2$-Saponine | 15 | 0.0 | 0.0 | 0.0 | 0.0 | 0.0 | 0.0 |
| HEK293T | FPV | 15 | 82.9 | 7.3 | 81.9 | 4.9 | 1.3 | 2.7 |
| HEK293T | WSN | 15 | 90.5 | 3.8 | 90.6 | 3.6 | 0.9 | 2.0 |
| DF1 | MOCK | 18 | 0.0 | 0.0 | 0.0 | 0.0 | 0.0 | 0.0 |
| DF1 | $H_2O_2$-Saponine | 15 | 0.0 | 0.0 | 0.0 | 0.0 | 0.0 | 0.0 |
| DF1 | FPV | 15 | 63.6 | 12.5 | 64.8 | 8.3 | 2.2 | 4.6 |
| DF1 | WSN | 15 | 84.6 | 20.2 | 83.1 | 14.7 | 3.8 | 8.1 |

**Table S2: Overview of the RG ratio values from the FRET analysis for HEK293T and DF1 cells.** Data correspond to Figure 1 of the main manuscript. n: number of cells, iqr: interquartile range, sd: standard deviation, se: standard error of the mean, ci: confidence interval.

| cell type | treatment | n | median | iqr | mean | sd | se | ci |
|---|---|---|---|---|---|---|---|---|
| HEK293T | MOCK | 50 | 0.21 | 0.05 | 0.22 | 0.03 | 0.00 | 0.01 |
| HEK293T | DOPS | 55 | 0.61 | 0.11 | 0.64 | 0.08 | 0.01 | 0.02 |
| HEK293T | FPV | 51 | 0.49 | 0.12 | 0.49 | 0.07 | 0.01 | 0.02 |
| HEK293T | WSN | 53 | 0.51 | 0.11 | 0.53 | 0.09 | 0.01 | 0.02 |
| DF1 | MOCK | 33 | 0.22 | 0.07 | 0.24 | 0.05 | 0.01 | 0.02 |
| DF1 | DOPS | 21 | 0.51 | 0.12 | 0.54 | 0.13 | 0.03 | 0.06 |
| DF1 | FPV | 33 | 0.47 | 0.15 | 0.50 | 0.12 | 0.02 | 0.04 |
| DF1 | WSN | 33 | 0.49 | 0.11 | 0.49 | 0.10 | 0.02 | 0.04 |

**Table S3: Overview of the GP index values from the analysis with Laurdan for HEK293T and DF1 cells.** Data correspond to Figure 2 of the main manuscript. n: number of cells, iqr: interquartile range, sd: standard deviation, se: standard error of the mean, ci: confidence interval.

| cell type | treatment | n | median | iqr | mean | sd | se | ci |
|---|---|---|---|---|---|---|---|---|
| HEK293T | MOCK | 52 | 0.01 | 0.12 | 0.00 | 0.09 | 0.01 | 0.03 |
| HEK293T | MbCD | 56 | -0.44 | 0.38 | -0.45 | 0.22 | 0.03 | 0.06 |
| HEK293T | FPV | 103 | 0.28 | 0.10 | 0.27 | 0.07 | 0.01 | 0.01 |
| HEK293T | WSN | 110 | 0.31 | 0.07 | 0.31 | 0.05 | 0.01 | 0.01 |
| DF1 | MOCK | 86 | -0.04 | 0.11 | -0.04 | 0.10 | 0.01 | 0.02 |
| DF1 | MbCD | 87 | -0.34 | 0.14 | -0.35 | 0.11 | 0.01 | 0.02 |
| DF1 | FPV | 61 | 0.25 | 0.15 | 0.25 | 0.10 | 0.01 | 0.03 |
| DF1 | WSN | 57 | 0.21 | 0.11 | 0.23 | 0.10 | 0.01 | 0.03 |

**Table S4: Overview of the GP index values from the analysis with Di-4-ANEPPDHQ for HEK293T and DF1 cells.** Data correspond to Figure 2 of the main manuscript. n: number of cells, iqr: interquartile range, sd: standard deviation, se: standard error of the mean, ci: confidence interval.

| cell type | treatment | n | median | iqr | mean | sd | se | ci |
|---|---|---|---|---|---|---|---|---|
| HEK293T | MOCK | 53 | 0.26 | 0.08 | 0.25 | 0.05 | 0.01 | 0.01 |
| HEK293T | MbCD | 53 | -0.01 | 0.08 | 0.01 | 0.08 | 0.01 | 0.02 |
| HEK293T | FPV | 116 | 0.44 | 0.03 | 0.44 | 0.02 | 0.00 | 0.00 |
| HEK293T | WSN | 127 | 0.44 | 0.03 | 0.44 | 0.02 | 0.00 | 0.00 |
| DF1 | MOCK | 36 | 0.31 | 0.04 | 0.31 | 0.03 | 0.01 | 0.01 |
| DF1 | MbCD | 76 | 0.07 | 0.12 | 0.07 | 0.08 | 0.01 | 0.02 |
| DF1 | FPV | 51 | 0.44 | 0.02 | 0.44 | 0.02 | 0.00 | 0.01 |
| DF1 | WSN | 63 | 0.44 | 0.04 | 0.44 | 0.02 | 0.00 | 0.01 |

**Table S5 Overview of the diffusion coefficient values [µm²/s] from the sFCS analysis for HEK293T and DF1 cells.** Data correspond to Figure 3 of the main manuscript. n: number of cells, iqr: interquartile range, sd: standard deviation, se: standard error of the mean, ci: confidence interval.

| protein | cell type | treatment | n | median | iqr | mean | sd | se | ci |
|---|---|---|---|---|---|---|---|---|---|
| mp-mEGFP | HEK293T | MOCK | 31 | 1.28 | 0.29 | 1.28 | 0.17 | 0.03 | 0.06 |
| mp-mEGFP | HEK293T | FPV | 20 | 0.55 | 0.29 | 0.55 | 0.26 | 0.06 | 0.12 |
| mp-mEGFP | HEK293T | WSN | 21 | 0.63 | 0.18 | 0.65 | 0.19 | 0.04 | 0.09 |
| GPI-mEGFP | HEK293T | MOCK | 29 | 1.04 | 0.34 | 1.06 | 0.18 | 0.03 | 0.07 |
| GPI-mEGFP | HEK293T | FPV | 20 | 0.64 | 0.25 | 0.63 | 0.23 | 0.05 | 0.11 |
| GPI-mEGFP | HEK293T | WSN | 25 | 0.66 | 0.34 | 0.61 | 0.22 | 0.04 | 0.09 |
| HA-mEGFP | HEK293T | MOCK | 36 | 0.42 | 0.14 | 0.44 | 0.11 | 0.02 | 0.04 |
| HA-mEGFP | HEK293T | FPV | 20 | 0.39 | 0.29 | 0.39 | 0.17 | 0.04 | 0.08 |
| HA-mEGFP | HEK293T | WSN | 30 | 0.33 | 0.12 | 0.35 | 0.14 | 0.03 | 0.05 |
| mp-mEGFP | DF1 | MOCK | 22 | 1.17 | 0.32 | 1.18 | 0.23 | 0.05 | 0.10 |
| mp-mEGFP | DF1 | FPV | 20 | 0.70 | 0.39 | 0.67 | 0.22 | 0.05 | 0.10 |
| mp-mEGFP | DF1 | WSN | 20 | 0.55 | 0.36 | 0.55 | 0.24 | 0.05 | 0.11 |
| GPI-mEGFP | DF1 | MOCK | 20 | 0.99 | 0.34 | 0.99 | 0.24 | 0.05 | 0.11 |
| GPI-mEGFP | DF1 | FPV | 18 | 0.57 | 0.19 | 0.55 | 0.15 | 0.04 | 0.08 |
| GPI-mEGFP | DF1 | WSN | 20 | 0.47 | 0.23 | 0.49 | 0.14 | 0.03 | 0.07 |
| HA-mEGFP | DF1 | MOCK | 18 | 0.44 | 0.17 | 0.48 | 0.11 | 0.03 | 0.06 |
| HA-mEGFP | DF1 | FPV | 18 | 0.27 | 0.15 | 0.33 | 0.11 | 0.03 | 0.05 |
| HA-mEGFP | DF1 | WSN | 22 | 0.29 | 0.11 | 0.29 | 0.09 | 0.02 | 0.04 |

#### 3. Figures

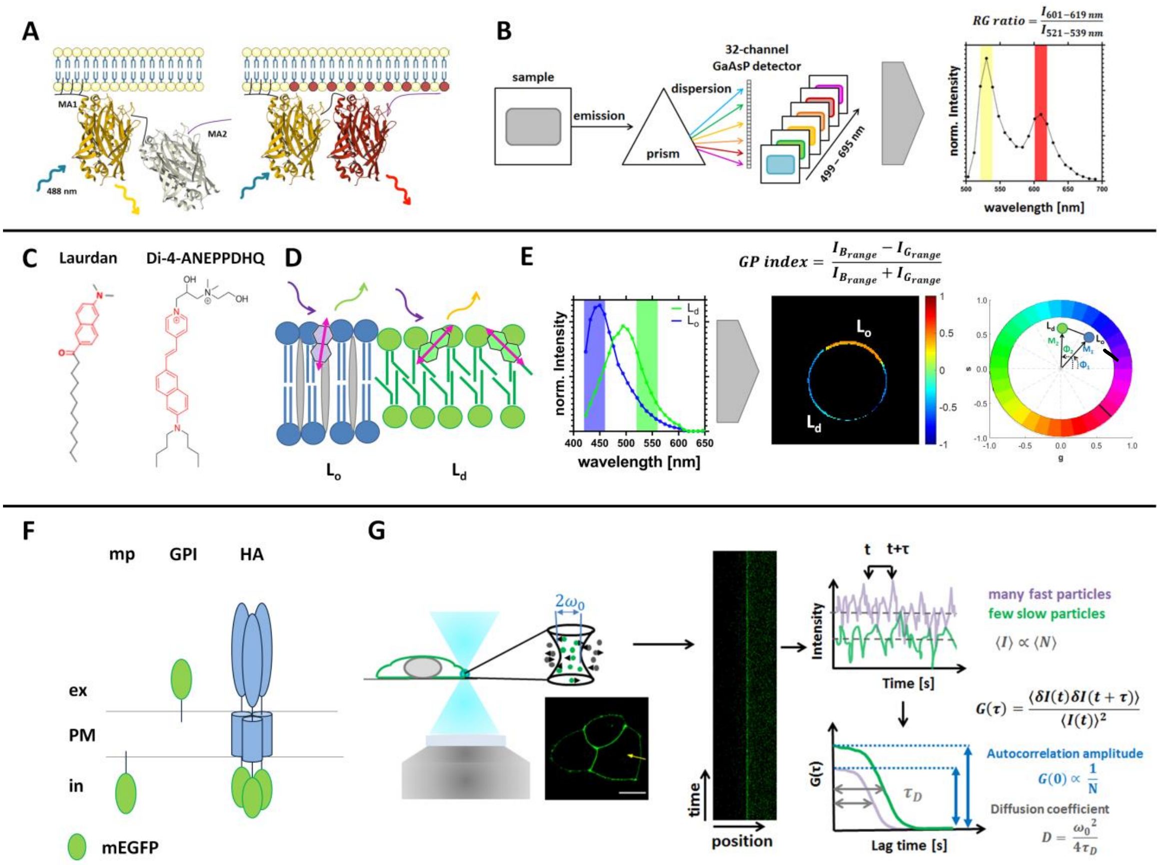


**Figure S1: Overview of experimental approaches used in this study.** Schematic overview of the experimental setup for the analysis of the enrichment of negatively charged lipids at the inner leaflet of the PM via FRET (A-B), changes in membrane fluidity via GP index (C-E), and dynamics of PM-associated proteins using sFCS (F-G). **(A)** Cartoon of the strategy used to study the effective electrostatic potential of the inner leaflet of the plasma membrane (PM). The lipid charge FRET-sensor (MCS+) contains two membrane attachment (MA) units and two fluorescent proteins (FPs, Venus and mCherry). MA1 is linked to the N-terminus of the FP Venus and consists of two palmitoylation and one myristoylation sites with the LCK10 sequence (MGCVCSSNPE), which allows a permanent binding to the PM via hydrophobic interactions. MA2 (purple) is linked to the C-terminus of the FP mCherry and contains a polybasic sequence (ARFGRRRRRRIRFRWVIM) which associates to negatively charged lipids at the inner leaflet of the PM. The FRET efficiency is then low when MA2 is separated from the PM (left panel) and high upon association of MA2 to the PM (right panel). Adapted from [4]. **(B)** Schematic illustration of the spectral FRET imaging analysis. The fluorescence emission from the sample upon illumination with laser light (488 nm) is dispersed (using a prism or other device) and guided onto a 32-channel GaAsP detector. Each channel records the signal at different wavelengths (499-695 nm), with a bin width of 8.9 nm. The signals are then used to visualize the emission spectrum for each pixel in the image. The intensity ratio (RG ratio) is then calculated from the spectrum values at each pixel (and averaged for all the pixels in a ROI and for different ROIs, if needed). Adapted from [5]. **(C)** Chemical structures of the solvatochromic probes Laurdan and Di-4-ANEPPDHQ used for the determination of the membrane fluidity at the PM, with the fluorophores highlighted in red. Parts in red highlight the fluorophore in the different probes. Taken from [6]. **(D)** Working principle of the solvatochromic probe Laurdan for lipid bilayers, presented as a mix of lipid ordered ($L_o$, blue, with cholesterol in grey) and lipid disordered ($L_d$, green) phases. Magenta arrows represent the orientation of Laurdan in the membrane. The interaction with the local environment leads to a spectral shift. The emission is shifted towards shorter wavelengths when the probe is in the $L_o$ phase and towards longer in the $L_d$ phase. Adapted from [7, 8]. **(E)** The emission signals of the spectral imaging (as shown in panel B) were then used to visualize the emission spectrum for each image. Representative fluorescence spectra for Laurdan in the $L_o$ (blue) and $L_d$ (green) phases of lipid bilayers. Intensity shifts between the $L_o$ and $L_d$ phases region can be quantified through a generalized polarization (GP) index.

**Figure S1 (continued):** GP value range from −1.0 (very fluid) to 1.0 (very gel-like) and are calculated for each image pixel in order to obtain a membrane "fluidity" map. The emission spectra are alternatively used to obtain a phasor plot. Spectral phasor plots represent the spectra as vectors of modulation (M) and phase angle (Φ), which are related to the spectral width and emission maximum ($\lambda_{max}$). The phase angle moves counterclockwise with increasing $\lambda_{max.}$ An increase in spectral width shifts the phasor closer to the center. Single dots represent exemplificative phasor values for a fluid membrane ($L_d$, green) and a rigid membrane ($L_o$, blue). Adapted from [7, 8]. **(F)** Overview of the different proteins used to study the dynamics of membrane components via sFCS. Monomeric membrane associated constructs consisting of a myristoylated and palmitoylated (mp) peptide or glycosylphosphatidylinositol (GPI)-anchor linked to the monomeric fluorescent protein mEGFP are used to probe the diffusion behavior in the inner and outer leaflet of the PM, respectively. As a model of transmembrane protein, we selected the trimeric hemagglutinin (HA) receptor from IAV, which is C-terminal linked to mEGFP. **(G)** Schematic principle of a confocal based sFCS setup. The focused laser beam scans the sample perpendicular to the PM (scan line is shown as kymograph), where mEGFP-linked proteins diffuse in and out of the confocal volume, giving rise to fluorescence fluctuations. From the resulting intensity trace, the autocorrelation function (G(τ)), which represents the self-similarity of the signal, is calculated and fitted to a two-dimensional diffusion model in order to obtain diffusion time ($\tau_D$, half-maximum decay of G(τ)) and the concentration of the diffusing particles (N, from the ACF value at time zero). An increase of $\tau_D$ is associated with slower diffusive dynamics and a decrease of the ACF amplitude corresponds to an increase of N. Adapted from [9].

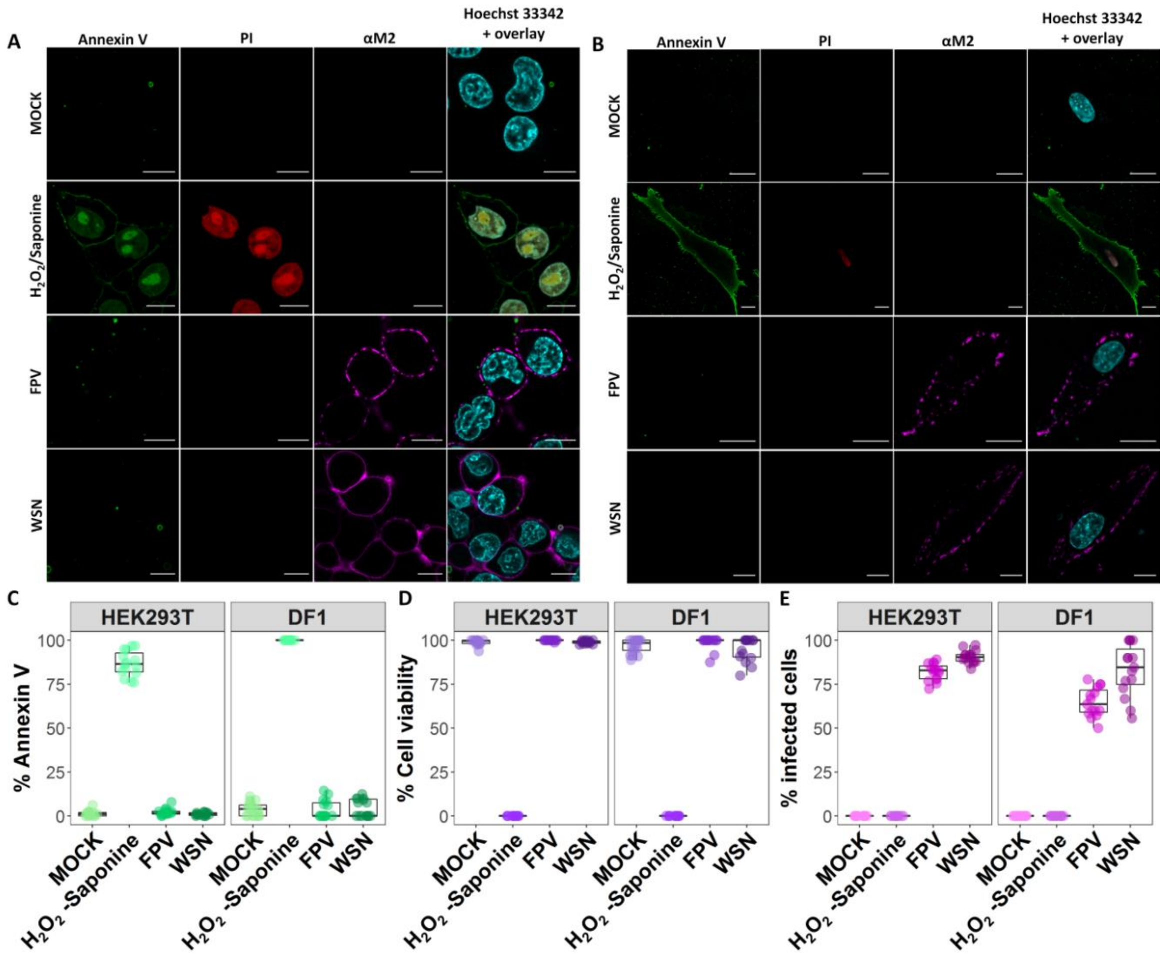


**Figure S2: Early-stage IAV infection does not impact cell viability and PS trans-leaflet organization in the investigated cell models.** HEK293T and DF1 cells were either: non-infected (MOCK), treated with 8 µM $H_2O_2$/0.1% Saponine (positive control), infected with FPV or infected with WSN. Samples were co-stained 16 hpi with Annexin V-AF488, propidium iodide (PI), anti-M2-AF647 (αM2) and Hoechst 33342. Representative confocal scanning microscope images of co-stained HEK293T cells **(A)** and DF1 cells **(B)**. Apoptotic cells exposing PS in the outer leaflet of the PM were visualized with Annexin V (green), dead/non-viable cells with PI (red), infected cells with αM2 (magenta) and the cell nucleus with Hoechst 33342 (cyan). Scale bars: 10 µm. Box plot with single data points of two independent experiments show the percentages of apoptotic (Annexin V-positive) cells **(C)**, the percentages of viable (Annexin V-/PI-negative) cells **(D)** and the percentage of infected (αM2-positive) cells **(E)**, which were calculated in relation to the total amount of cells (Hoechst 33342-positive). Quantitative information and statistical description are summarized in Table S1. For each condition, 15 to 18 images were manually analyzed, including in total more than 830 HEK293T cells and more than 150 DF1 cells per treatment.-

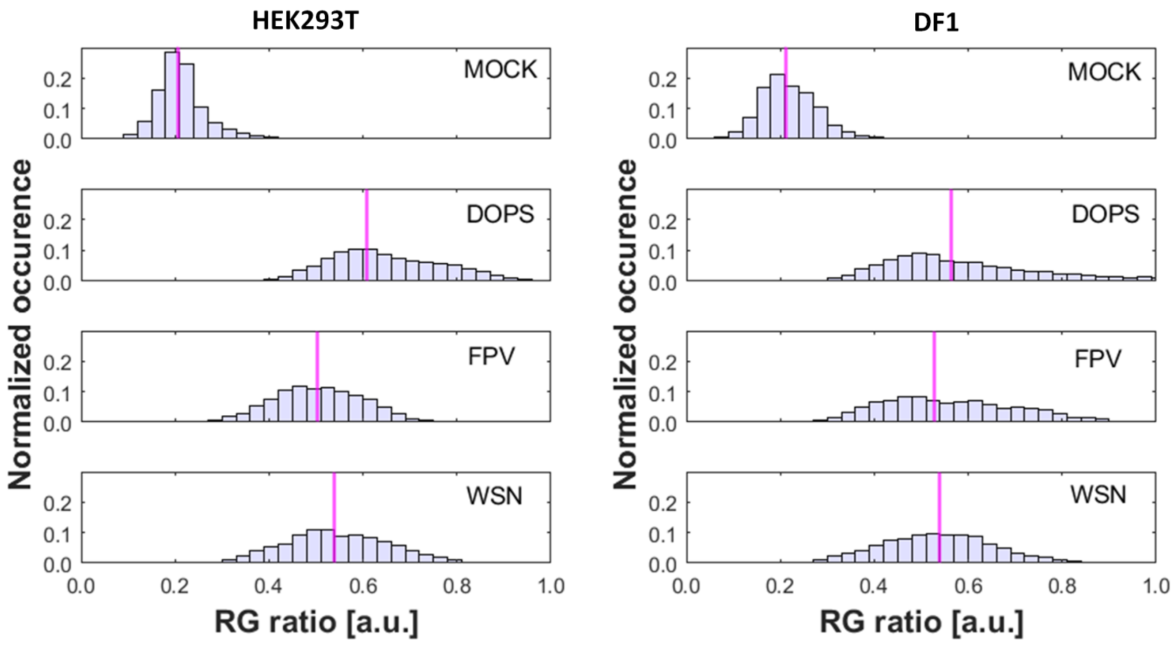


**Figure S3: Quantification of the RG ratio from FRET measurements at the PM of HEK293T and DF1 cells.** HEK293T and DF1 cells were either: non-infected (MOCK), treated with DOPS-SUVs (positive control), infected with FPV or WSN. All cells were expressing the FRET-sensor MCS+ and emission spectrum images were acquired 16 hpi. The obtained RG ratio values from each pixel in all ROIs selected at the PM of 50-55 HEK293T cells and 21-33 DF1 cells (Table S2) for all conditions were pooled and represented as normalized histogram showing the median value (magenta). Data correspond to the analyzed cells in Figure 1 of the main manuscript.

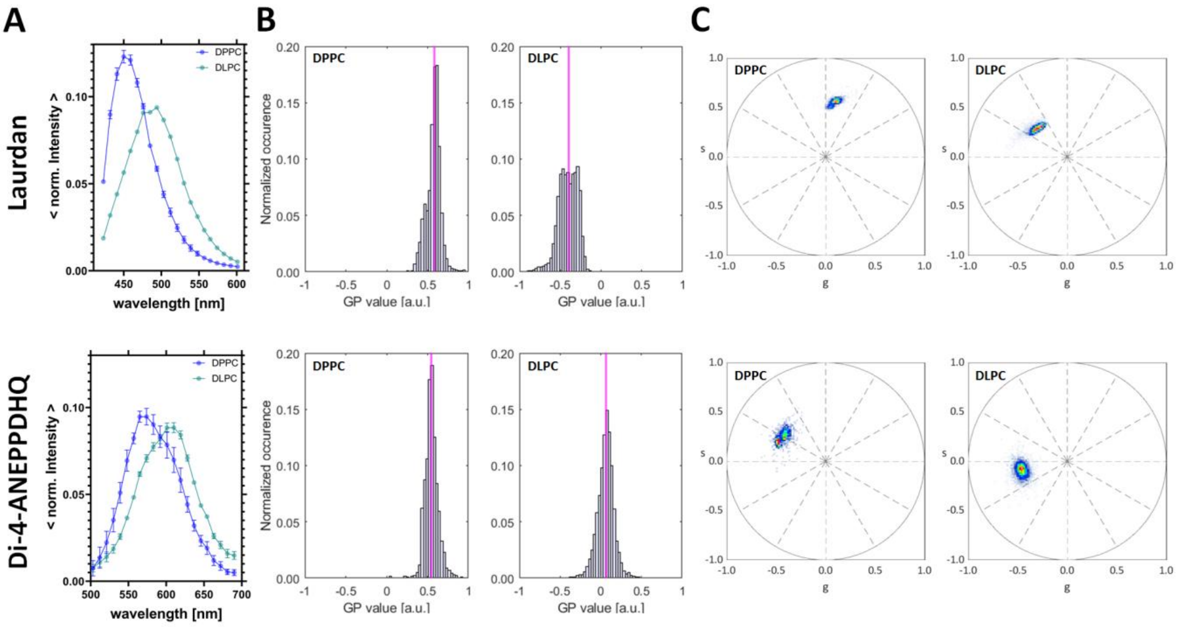


**Figure S4: GP and spectral phasor analysis of GUVs labelled with Laurdan and Di-4-ANEPPDHQ.** In order to acquire reference points for the GP and phasor analysis of infected cells, GUVs with extremely different lipid bilayer environments (solid ordered DPPC and liquid disordered DLPC) were prepared. These samples were then stained with Laurdan or Di-4-ANEPPDHQ and imaged with the same acquisition parameters used for the analysis of infected cells. For this analysis, 15 DLPC GUVs and 10 DPPC GUVs were analyzed for each dye. **(A)** Averaged normalized intensity spectra of all selected ROIs in the two types of GUVs. Data are represented as mean ± SD. **(B)** The obtained pixel-wise GP values of all ROIs GUVs were pooled and represented as normalized histogram, with the median value highlighted in magenta. The median values for DPPC were 0.58 (Laurdan) and 0.55 (Di-4-ANEPPDHQ). For DLPC they were -0.40 (Laurdan) and 0.07 (Di-4-ANEPPDQ). **(C)** Spectral phasor plots are obtained as described in Paragraph 1.3. Each point in the plot corresponds to s and g coordinates calculated from the pixel-wise spectrum of each ROI in a GUV. In the presence of a more ordered lipid environment, the point cloud shifts “clockwise” in the plot.

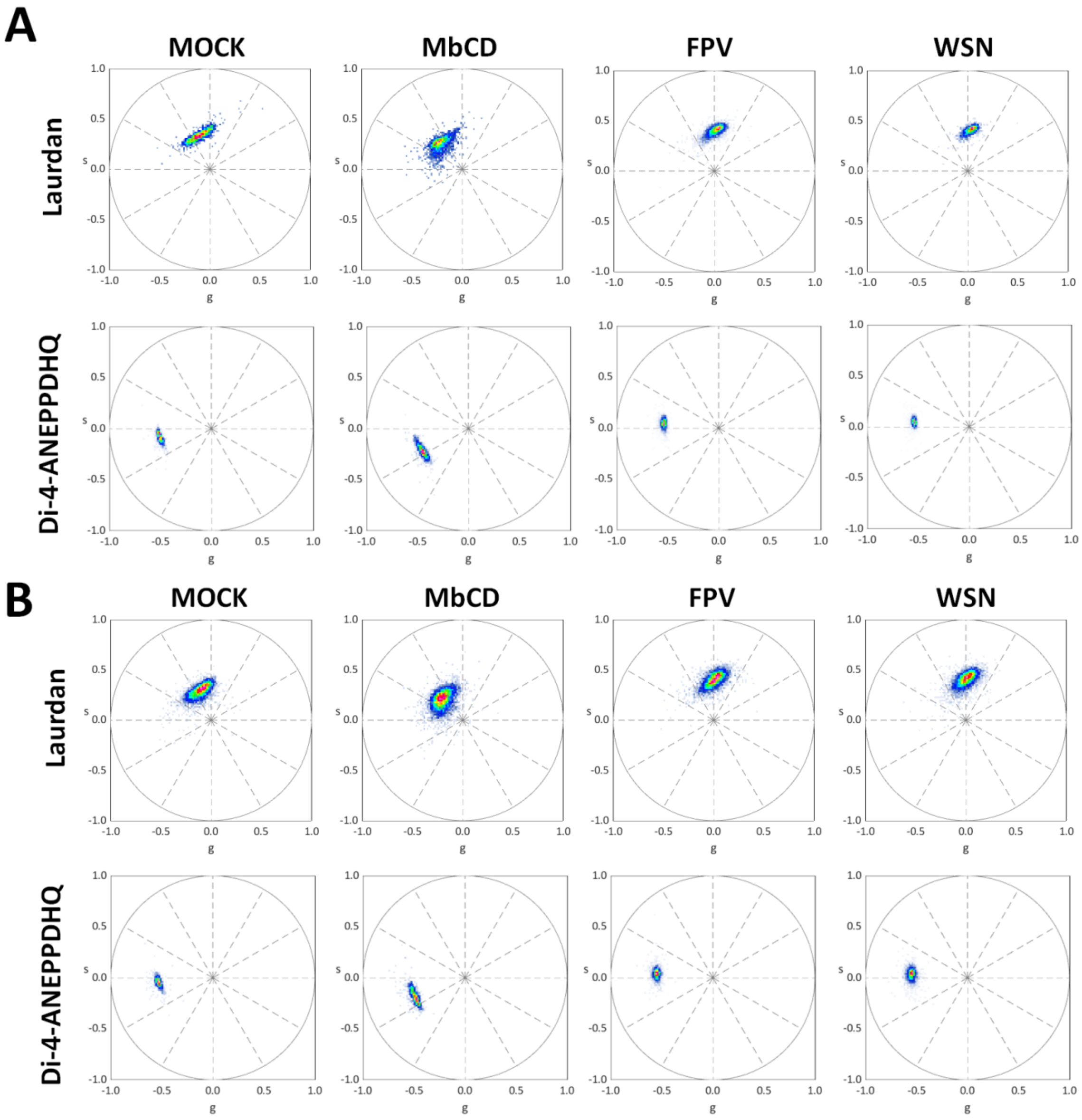


**Figure S5: Spectral phasor analysis of Laurdan- and Di-4-ANEPPDHQ-labelled HEK293T and DF1 cells.** HEK293T **(A)** and DF1 **(B)** cells were either: non-infected (MOCK), treated with methyl-β-cyclodextrin (MbCD), infected with FPV or infected with WSN. All cells were labelled with Laurdan and Di-4-ANEPPDHQ and emission spectrum images were acquired 16 hpi. For this analysis, 52-110 Laurdan-stained cells and 36-127 Di-4-ANEPPDHQ-stained cells were selected (Table S3 and S4) Spectral phasor plots are obtained as described in Paragraph 1.3. Each point in the plot corresponds to s and g coordinates calculated from the pixel-wise spectrum of each selected ROI. Data correspond to the analyzed cells in Figure 2 of the main manuscript.

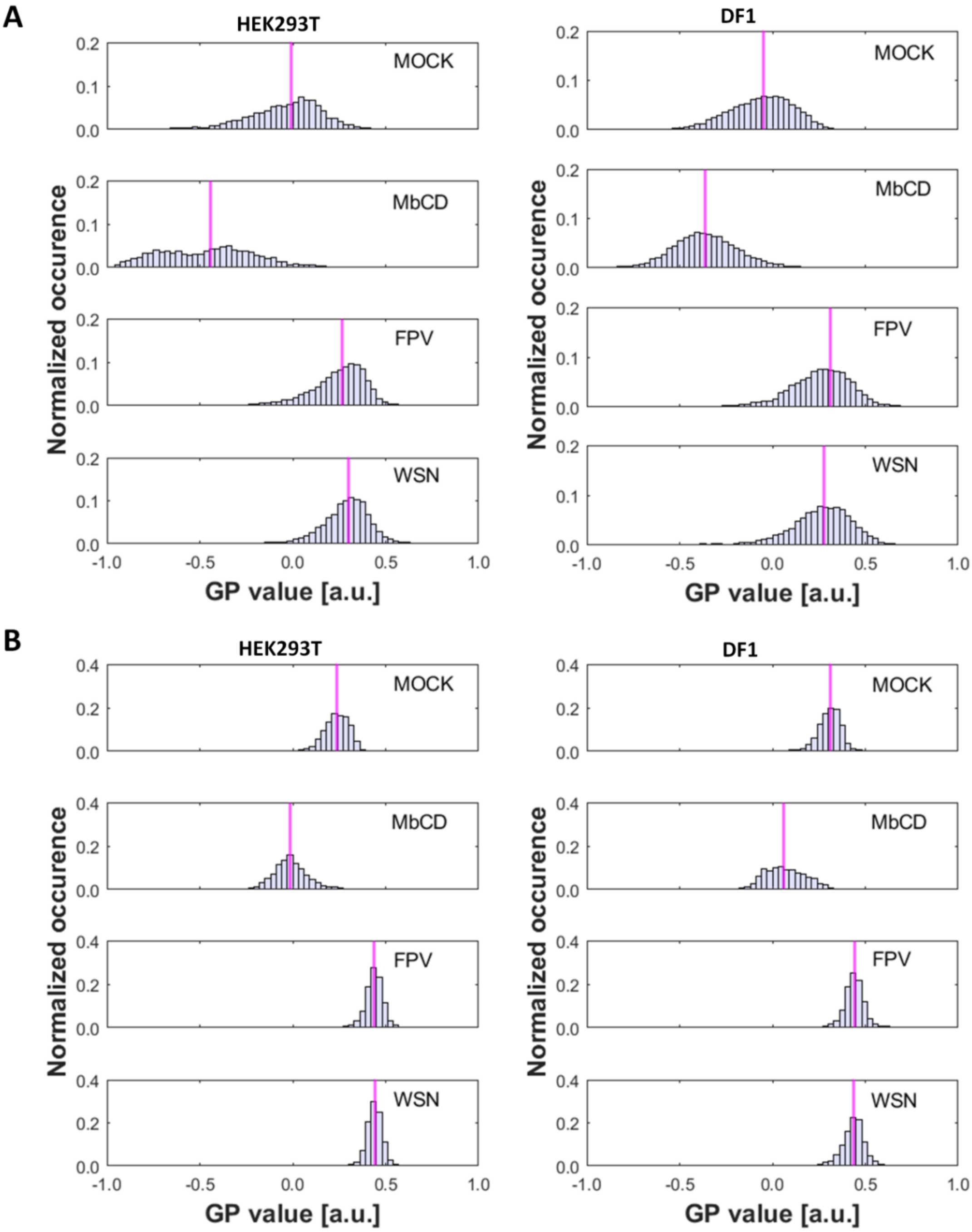


**Figure S6: Quantification of the GP index from membrane fluidity measurements at the PM of HEK293T and DF1 cells.** HEK293T and DF1 cells were either: non-infected (MOCK), treated with methyl-β-cyclodextrin (MbCD), infected with FPV or infected with WSN. All cells were labelled with Laurdan **(A)** and Di-4-ANEPPDHQ **(B)** and emission spectrum images were acquired 16 hpi. For this analysis, 52-110 Laurdan stained cells and 36-127 Di-4-ANEPPDHQ cells were selected (Table S3 and S4). Obtained pixel-wise GP values from all selected PM ROIs were pooled for each condition and represented as normalized histogram showing the median value (magenta). Descriptive statistics are summarized in Table S3-S4.Data correspond to the analyzed cells in Figure 2 of the main manuscript.

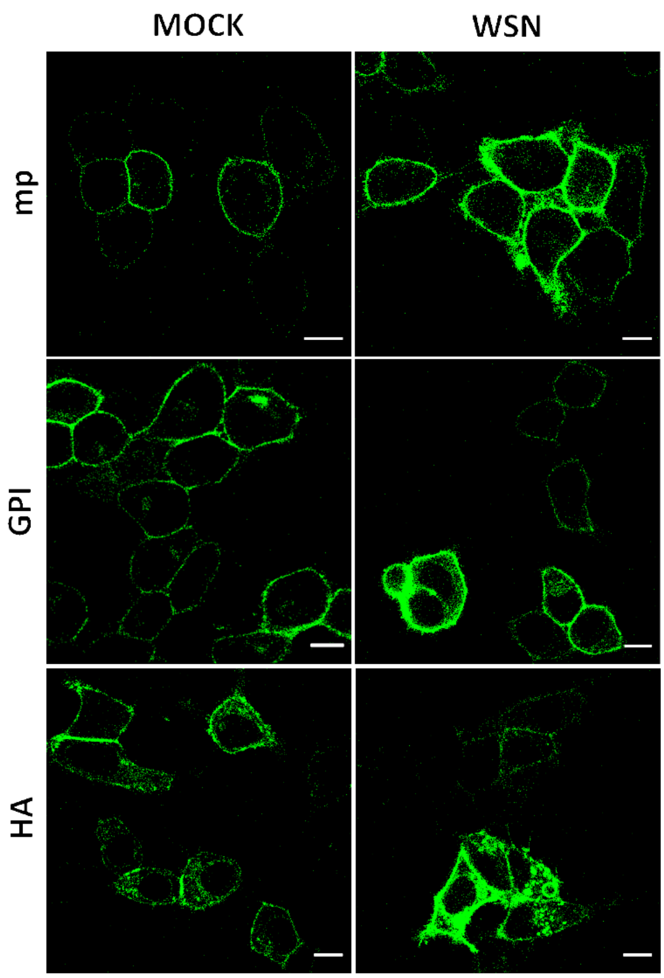


**Figure S7: Representative confocal fluorescence images of HEK293T cells expressing mEGFP-tagged proteins.** HEK293T cells were transfected with mp-mEGFP, GPI-mEGFP or HA-mEGFP. After four hours, cells were infected with WSN or treated with simple medium (MOCK). The images acquired after 16 hours indicate no major differences in the fluorescence protein appearance between infected and non-infected cells. Cells with very high expression levels could be seen more often following infection, but those cells were not selected for further analysis. Cells with expression levels similar to those observed in MOCK samples were selected instead. Similar results were obtained for cells infected with FPV (data not shown). Scale bars represent 10 μm.

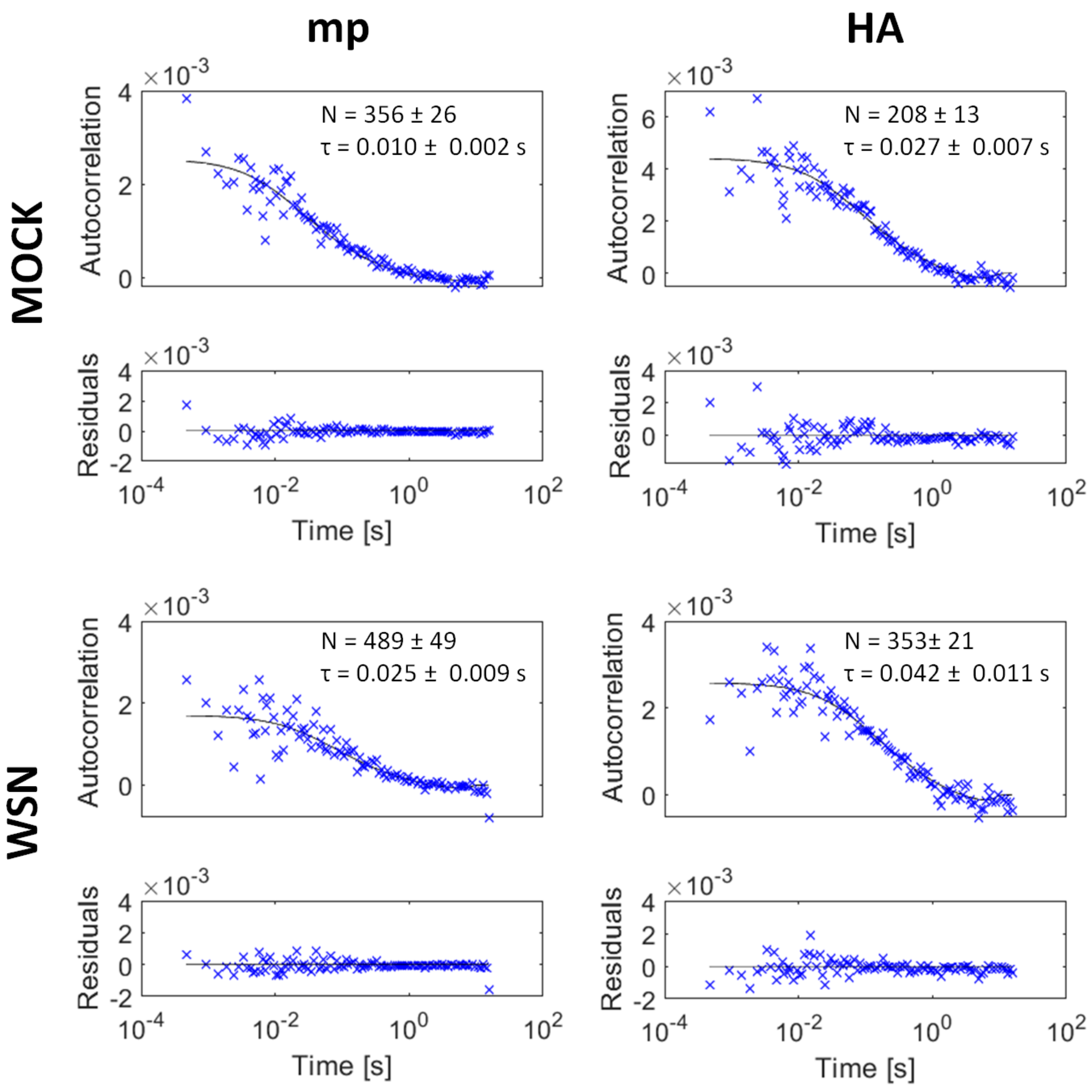


**Figure S8: Examples of autocorrelation functions.** Representative sFCS autocorrelation functions and fit curves obtained for non-infected (MOCK) and WSN-infected HEK293T cells expressing mp-mEGFP and HA-mEGFP. Fit curves (solid line) were obtained by fitting a two-dimensional diffusion model to the data, as described in the main text (Methods section). Data correspond to Figure 3 of the main manuscript. Similar data were obtained for DF1 cells (data not shown).
